# Mechanosensitive Expression of Lamellipodin Governs Cell Cycle Progression and Intracellular Stiffness

**DOI:** 10.1101/2020.10.30.362624

**Authors:** Joseph A. Brazzo, Kwonmoo Lee, Yongho Bae

## Abstract

Cells exhibit pathological behaviors in response to increased extracellular matrix (ECM) stiffness, including accelerated cell proliferation and migration [1–9], which are correlated with increased intracellular stiffness and tension [2, 3, 10–12]. The biomechanical signal transduction of ECM stiffness into relevant molecular signals and resultant cellular processes is mediated through multiple proteins associated with the actin cytoskeleton in lamellipodia [2, 3, 10, 11, 13]. However, the molecular mechanisms by which lamellipodial dynamics regulate cellular responses to ECM stiffening remain unclear. Previous work described that lamellipodin, a phosphoinositide- and actin filament-binding protein that is known mostly for controlling cell migration [14–21], promotes ECM stiffness-mediated early cell cycle progression [2], revealing a potential commonality between the mechanisms controlling stiffness-dependent cell migration and those controlling cell proliferation. However, i) whether and how ECM stiffness affects the levels of lamellipodin expression and ii) whether stiffness-mediated lamellipodin expression is required throughout cell cycle progression and for intracellular stiffness have not been explored. Here, we show that the levels of lamellipodin expression in cells are significantly increased by a stiff ECM and that this stiffness-mediated lamellipodin upregulation persistently stimulates cell cycle progression and intracellular stiffness throughout the cell cycle, from the early G1 phase to M phase. Finally, we show that both Rac activation and intracellular stiffening are required for the mechanosensitive induction of lamellipodin. More specifically, inhibiting Rac1 activation in cells on stiff ECM reduces the levels of lamellipodin expression, and this effect is reversed by the overexpression of activated Rac1 in cells on soft ECM. We thus propose that lamellipodin is a critical molecular lynchpin in the control of mechanosensitive cell cycle progression and intracellular stiffness.

## RESULTS AND DISCUSSION

### ECM stiffness regulates the levels of lamellipodin and cell cycle-associated mRNAs and proteins

ECM stiffness affects the expression and activation of critical effector proteins vital to ECM stiffness-dependent cellular functions, including the cell cycle and proliferation [1–5]. Our previous study provided the first evidence that lamellipodin is a key mediator that transduces ECM stiffness into early cell cycle progression (at the G1 phase) of mouse embryonic fibroblasts (MEFs) [2]. However, whether the expression of lamellipodin is regulated by ECM stiffness has never been investigated. To determine whether the levels of lamellipodin mRNA and protein expression are stiffness-dependent, we used fibronectin-coated polyacrylamide hydrogels to create culture matrices that mimic the biologically relevant stiffness of normal and pathological arterial microenvironments *in vivo*, which are much less stiff than glass or plastic. Hydrogels of low stiffness (2-4 kPa; hereafter called “soft”), intermediate stiffness (10-12 kPa; hereafter called “medium”), and high stiffness (20-24 kPa; hereafter called “stiff”) stiffness approximate the normal and pathological stiffness of mouse arteries [1, 2]. This *in vitro* hydrogel system has been used for molecular and biomechanical analyses of cellular function in response to a given biomechanical stimulus (stiffness) by ECM [1–12]. Using this system (Figure 1A), serum-starved MEFs were cultured on fibronectin-coated hydrogels with 10% fetal bovine serum (FBS) for 9 and 24 hours. First, we found that the levels of lamellipodin mRNA expression, as determined by RT-qPCR, to be moderately increased at both 9 and 24 hours on stiff hydrogels (Figure 1B). We also confirmed that the mRNA expression levels of cyclin A (an S/G2-phase marker, Figure 1C) and cyclin D1 (an early G1 phase marker, Figure 1D) are stiffness-sensitive, as previously shown in other studies [1, 2]. Using immunoblotting, we then observed that the protein levels of lamellipodin in MEFs on stiff hydrogels were sustained for 9 (Figure 1F) and 24 (Figure 1G) hours, with a greater increase in lamellipodin expression at 24 hours. In addition, we demonstrated that upregulated protein levels of cyclin A and cyclin D1 depended on both time and ECM stiffness.

**Figure 1.**
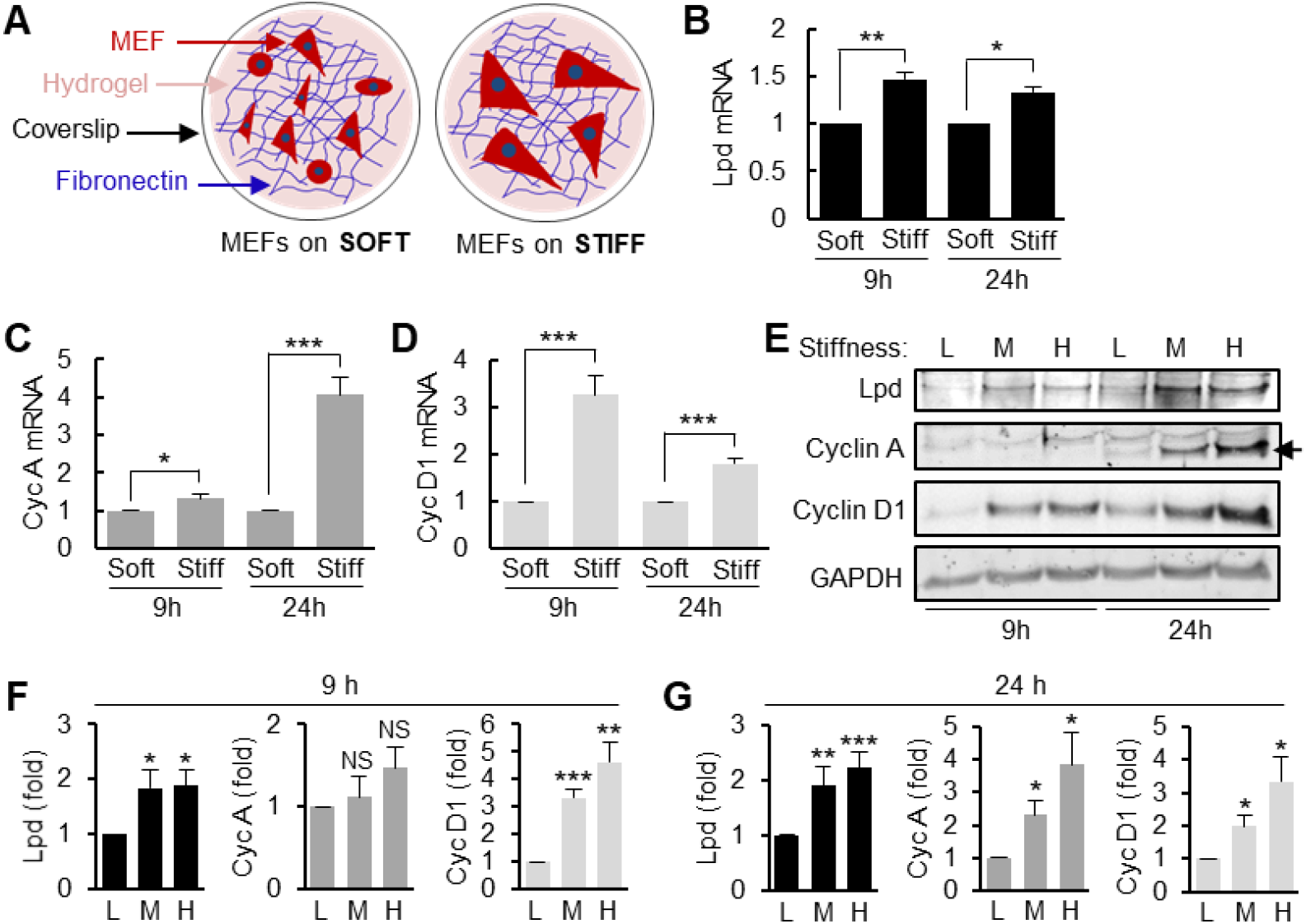
Stiff ECM increases the level of lamellipodin, cyclin A, and cyclin D1. **(A)** An *in vitro* model of tissue stiffening. Serum-starved MEFs were seeded on fibronectin-coated soft, medium, or stiff (H; 20-24 kPa) hydrogels and cultured with 10% FBS for 9 and 24 h. mRNAs **(B-D)** and total cell lysates **(E)** were analyzed by RT-qPCR and immunoblotting, respectively. Levels of lamellipodin (Lpd, **B**), cyclin A (Cyc A, **C**), and cyclin D1 (Cyc D1, **D**) mRNA were normalized to MEFs on soft hydrogels. 3 to 5 independent biological replicates. **(E)** A representative immunoblot image of 3 to 5 independent biological replicates. An arrow represents the position of cyclin A protein. **(F-G)** The bar graphs show lamellipodin, cyclin A, and cyclin D1 protein levels at 9 h **(F)** and 24 h **(G)** normalized to MEFs on soft hydrogels, respectively. Error bars in C-E and F-G show mean + SEM. *p<0.05, **p<0.01, ***p<0.001. NS: not significant.

How lamellipodin and cyclins directly interact in response to ECM stiffness will provide further insight into the coupling of stiffness-sensitive proliferation. Taken together, our results show that lamellipodin is upregulated by stiff ECM, and its stiffness-sensitive expression may be associated with cell cycle progression.

### Lamellipodin is required for persistent stimulation of cell cycle progression and intracellular stiffening throughout the entire cell cycle

Lyulcheva *et al*. showed, for the first time, that pico (its mammalian ortholog, lamellipodin) expressed in *Drosophila* and lamellipodin expressed in HeLa cells cultured on rigid tissue culture plates are required to promote cell proliferation [22]. Interestingly, we previously showed that lamellipodin transduces ECM stiffness into early cell cycle progression by upregulating the levels of cyclin D1 induction in MEFs cultured on stiff hydrogels [2]. However, to our knowledge, the degree to which lamellipodin mediates mechanosensitive cell cycle progression (or specific cell cycle stages) that contributes to cell proliferation remains unknown. To determine the broadscale effect of lamellipodin on cell cycle progression, we first sought to explore cell-cycle transcriptional networks regulated by lamellipodin using Ingenuity Pathway Analysis [23]. We found an interactive gene network associated with lamellipodin (RAPH1), cyclin A (or CCNA1), and cyclin D (or CCND1) (Figure 2A). We then determined the ability of mechanosensitive lamellipodin in MEFs to affect specific cell cycle stages. MEFs were transfected with small interfering RNA (siRNA) to acutely and transiently knockdown lamellipodin (Figure 2B and D). Interestingly, knocking down lamellipodin expression significantly reduced cyclin A (S/G2-phase) and cyclin B (G2/M-phase) (Figure 2B-C) when MEFs were cultured on stiff hydrogels for 24 hours. We additionally confirmed that cyclin D1 (early G1 phase) induction at 9 hours was markedly decreased by lamellipodin knockdown (Figure 2D-E), as previously shown [2]. Collectively, our results reveal that lamellipodin is essential for sustained mechanosensitive cell cycle progression.

**Figure 2.**
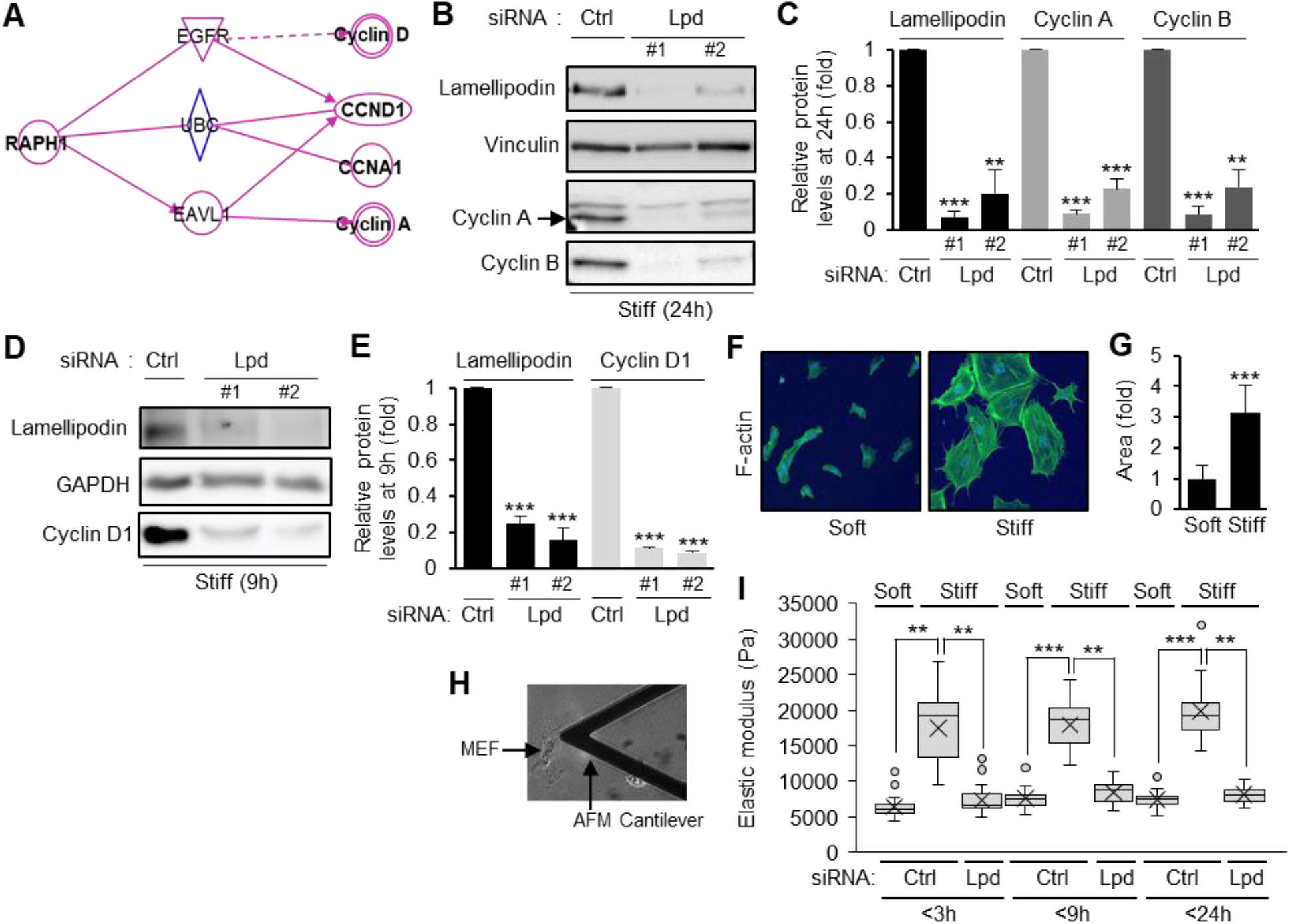
Loss of lamellipodin expression significantly inhibits cell cycle progression and intracellular stiffness. **(A)** The Ingenuity Pathway Analysis [23] Path Explorer tool was used to identify experimentally observed direct or indirect signaling intermediates downstream of lamellipodin *(RAPH1;* Ras Association (RalGDS/AF-6) And Pleckstrin Homology Domains 1). Symbol codes can be found at https://digitalinsights.qiagen.com/support/tutorials/. **(B-E)** MEFs were transfected with control (ctrl) siRNA or siRNAs to lamellipodin (Lpd #1 and Lpd #2), serum-starved, and seeded onto stiff hydrogels with 10% FBS for 24 h **(B)** or 9 h **(D)**, respectively. Total cell lysates were analyzed by immunoblotting. The bar graph shows lamellipodin, cyclin A, and cyclin B **(C)** or lamellipodin and cyclin D1 **(E)**, respectively. The protein levels were normalized to MEFs treated with control siRNA. Images in C and E are representative of n=3 independent biological replicates. Error bars in C and E show mean + SEM. **p<0.01, ***p<0.001. **(F)** Fixed MEFs cultured on soft and stiff hydrogels were incubated with FITC-conjugated phalloidin to stain F-actin. **(G)** The area of cell spreading was analyzed using ImageJ and normalized to MEFs on soft hydrogels. 21 cells on soft hydrogels and 19 MEFs on stiff hydrogels were quantified. ***p<0.001. **(H)** A bright-field image showing AFM cantilever position on MEF. **(I)** MEFs transfected with ctrl or lamellipodin (Lpd) siRNAs were serum-starved and seeded on FN-coated soft or stiff hydrogels with 10% FBS as indicated. Intracellular stiffness (elastic modulus) was measured by AFM and summarized as a box and whisker plot. 23 cells were analyzed from three independent experiments. **p<0.01, ***p<0.001.

Cells modulate their intracellular stiffness in response to the stiffness of the surrounding ECM [10, 11, 24], and this intracellular stiffness is critical for cell cycle progression and thus for proliferation [1–3]. More specifically, intracellular stiffening is the dynamic result of spatiotemporally organized actin recruitment and resultant filamentous actin (F-actin) formation. Moreover, intracellular stiffening is greatest at the cell periphery [2, 25], where lamellipodin is an essential mediator of leading-edge formation and a regulator of F-actin recruitment and formation at the cell periphery [14–16, 20, 21]. Using phalloidin immunofluorescence, we confirmed that F-actin in MEFs on stiff hydrogels is mostly located at the cell periphery (Figure 2F), and these MEFs also exhibit a large increase in cell spreading compared to MEFs cultured on soft hydrogels (Figure 2G). We further tested whether knocking down lamellipodin expression affects ECM stiffness-mediated intracellular stiffness associated with cell cycle progression. Atomic force microscopy (AFM) was used to measure the stiffness of MEFs on soft and stiff hydrogels. For this experiment, serum-starved MEFs were seeded on fibronectin-coated hydrogels of both low and high stiffness and serum-stimulated with 10% FBS for 3, 9, and 24 hours. Our AFM data showed that stiff ECM increased the intracellular stiffness of MEFs (Figure 2I; columns 2, 5, and 8), suggesting that this increase in intracellular stiffness on stiff hydrogels may be the result of upregulated lamellipodin expression. Indeed, AFM analysis with MEFs showed that lamellipodin knockdown significantly reduced intracellular stiffness at 3 and 9 hours (Figure 2I; columns 3 and 6). We also confirmed the previously reported reduction in intracellular stiffening as a result of loss of lamellipodin expression at 24 hours [2] (Figure 2I; column 9). Taken together, these AFM results suggest that ECM stiffness-dependent lamellipodin expression constitutes a discrete signaling module that transduces sustained progression and coupling of mechanosensitive cell cycle progression and the associated intracellular stiffening.

### Rac1 and intracellular stiffening are required for mechanosensitive lamellipodin induction

A FAK-Rac1 signaling pathway is critical for the transmission of ECM stiffness into mechanochemical cues and the resultant changes to cell cycle progression, proliferation, and intracellular stiffness [1–3]. Upon the initial activation of this pathway by ECM stiffness, FAK activates the downstream effector Rac1, which was previously shown to directly bind to lamellipodin [16]. Activated Rac1 subsequently stimulates stiffness-dependent intracellular stiffening and early cell cycle progression through lamellipodin [2]. Although our results and those of a previous study showed that lamellipodin expression is sensitive to ECM stiffness (Figure 1B and E) and that Rac overexpression induces the translocation of lamellipodin to the cell periphery of MEFs plated on stiff hydrogels [2], whether the total level of lamellipodin expression is regulated by Rac1-mediated signaling remains to be explored. To further dissect the regulatory pathway by which ECM stiffness is translated into lamellipodin induction through stiffness-sensitive Rac activation, we first confirmed that Rac is persistently activated in MEFs in response to increased ECM stiffness at 24 hours (Figure 3A), where we observed significantly increased levels of lamellipodin, cyclin D1, and cyclin A mRNA and protein induction (Figure 1E and G). Since cell cycle progression in MEFs occurs over 24 hours, we expected persistent Rac1 activation (Figure 3A) and intracellular stiffening (Figure 2I) to be necessary for lamellipodin-mediated cell cycle progression. We then tested whether Rac1 is a potent upstream regulator of stiffness-dependent lamellipodin expression. For this experiment, MEFs treated with Rac1 siRNAs or adenovirus encoding Rac^N17^ (a dominant-negative Rac) were plated on fibronectin-coated stiff hydrogels and stimulated with 10% FBS for a total of 24 hours. The immunoblot analysis data showed that both siRNA- or adenovirus-mediated Rac1 inhibition significantly decreased lamellipodin induction (Figure 3B-E). In contrast, the overexpression of constitutively activated Rac (adenovirus-encoded Rac^V12^) dramatically rescued lamellipodin induction in MEF cultures on soft hydrogels for 24 hours (Figure 3F-G; columns 1 and 2).

**Figure 3.**
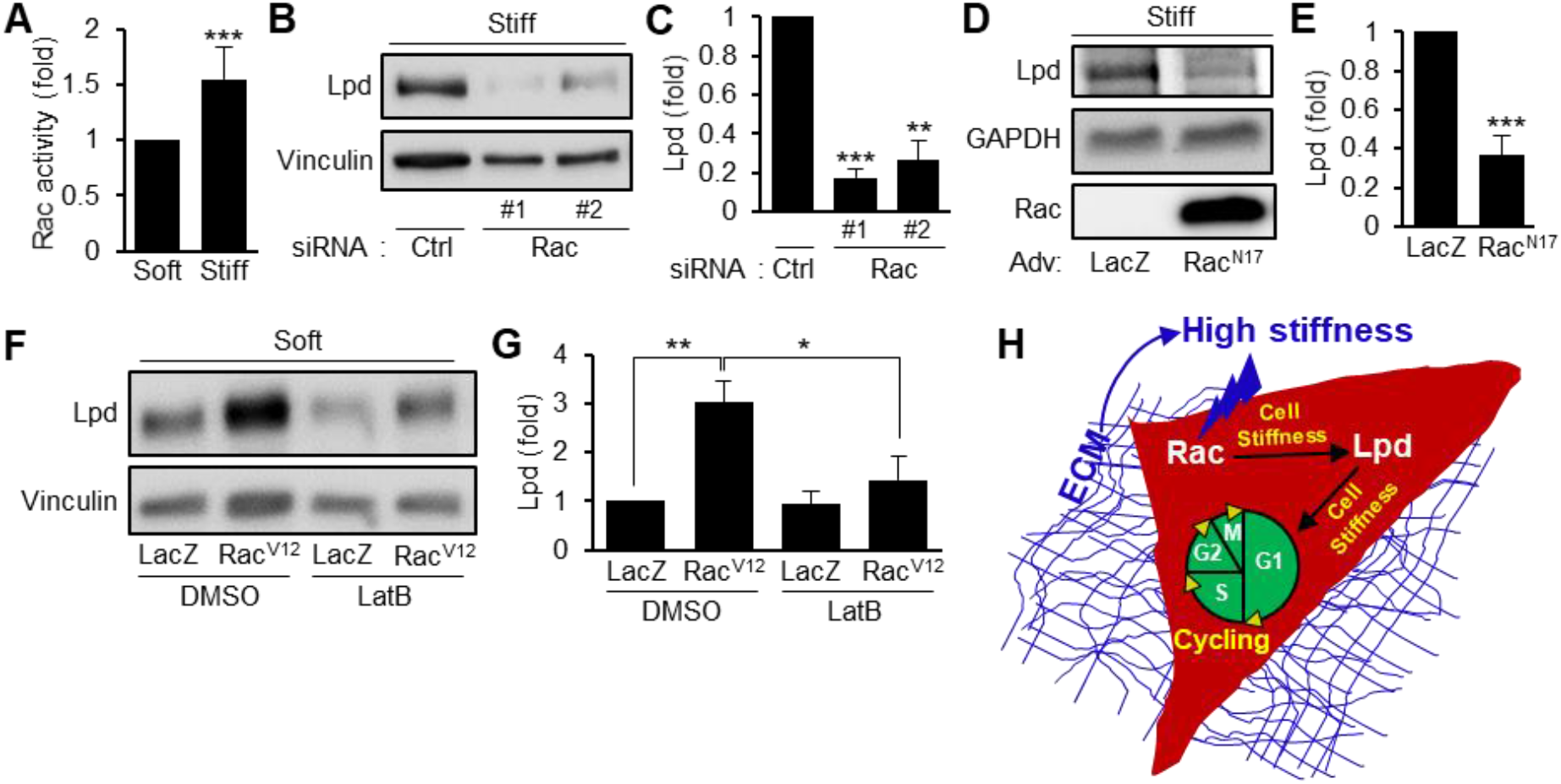
Rac and intracellular stiffening are required for mechanosensitive lamellipodin induction. (A) Serum-starved MEFs were cultured on soft or stiff hydrogels with 10% FBS for 24 hours. Rac activity was determined by G-LISA. *n=3* independent biological replicates. MEFs transfected with control or Rac1 siRNAs [26 and #2] (B), or infected with adenoviruses encoding LacZ or Rac^N17^(D), respectively, were seeded on FN-coated stiff hydrogels with 10% FBS for up to 24 hours and analyzed by immunoblotting. (C and E) The bar graphs show lamellipodin protein levels normalized to MEFs treated with control siRNA (B) or LacZ (D). (F) Immunoblots of MEFs infected with adenoviruses encoding LacZ or Rac^V12^ and seeded on soft hydrogels with 10% FBS ± latrunculin B (LatB) for 24 hours. (G) The bar graph shows lamellipodin protein levels normalized to MEFs infected with LacZ (DMSO). Error bars in A, C, E, and G show mean + SEM. *n=3* (B and F) and *n=4* (D) independent biological replicates. *p<0.05, **p<0.01, and ***p<0.001. (H) A model of mechanotransduction through the Rac-lamellipodin signaling module.

Intracellular forces (i.e., stiffness and tension) are generated by actin polymerization, which is largely regulated by Rac activity [2, 3]. Thus, we tested whether both Rac activation and intracellular stiffening are required for the increase in lamellipodin induction. For this experiment, MEFs infected with Rac^V12^ adenovirus were seeded on soft hydrogels with either DMSO (vehicle control) or latrunculin B (an actin polymerization inhibitor, which was previously shown to significantly reduce intracellular stiffness (Bae, 2014 #91) for 24 hours. Interestingly, we found that the overexpression of Rac1 in MEFs on the soft hydrogels was not sufficient to completely rescue Rac-mediated lamellipodin expression upon the decrease in intracellular stiffness [2] induced by latrunculin B treatment (Figure 3F-G, column 4), compared to the levels of lamellipodin expression in the Rac^V12^-infected cells treated with DMSO (Figure 3F-G, column 2). Considering these findings, we conclude that both Rac activation and intracellular stiffening are necessary for the mechanosensitive induction of lamellipodin (Figure 3H).

## CONCLUSIONS

Lamellipodin was first identified as an actin-associated Ena/VASP binding protein [14], and subsequent studies have shown lamellipodin to be essential for cell protrusion and migration [15–21]. However, i) the potential role of lamellipodin in governing other cellular functions, specifically cell proliferation and critically coupled cellular mechanical properties, and ii) the molecular pathway controlling lamellipodin expression in the context of mechanobiology have remained unexplored until now. Collectively, our results show that stable lamellipodin expression mediated by activated Rac1 is necessary for ECM stiffness-dependent sustained cell cycle progression and intracellular stiffening. Here, we shed new light on the mechanistic duality of lamellipodin in regulating ECM stiffness-dependent migration and cell cycle progression and intracellular stiffening, and we advance the exploration of lamellipodin as a molecular lynchpin useful for critically regulating lesser-known ECM stiffness-sensitive cellular functions.

## EXPERIMENTAL MODEL AND SUBJECT DETAILS

Spontaneously immortalized mouse embryonic fibroblasts (MEFs, a gift from the Assoian Laboratory, University of Pennsylvania) were grown in DMEM (Corning) supplemented with 50 μg/ml gentamicin solution (Corning) and 10% (v/v) fetal bovine serum (FBS, Sigma). Prior to seeding MEFs onto fibronectin-coated polyacrylamide hydrogels for experimentation, MEFs were synchronized in the G0 cell cycle phase (quiescence), in which near-confluent cells were serum-starved by incubation for 48 hours in DMEM with 1 mg/ml heat-inactivated, fatty-acid-free bovine serum albumin (BSA, TOCRIS). Thereafter, serum-starved MEFs were seeded onto fibronectin-coated polyacrylamide hydrogels with 10% FBS and incubated for 3, 9, and 24 hours.

## METHOD DETAILS

### siRNA transfection, adenovirus infection, and drug treatment

Near-confluent (80-90%) MEFs were transfected with 150 nM lamellipodin or Rac1 siRNAs using Lipofectamine 2000 reagent (Invitrogen) in reduced serum Opti-MEM as previously described [1, 2, 27].MEFs were transfected with siRNA for 5 hours, and then, approximately 48 hours after serum-starvation in serum-free DMEM with 1 mg/ml BSA, cells were trypsinized and reseeded onto fibronectin-coated hydrogels with fresh DMEM containing 10% FBS, as described above. A non-targeting scrambled siRNA (Silencer negative control siRNA, Ambion) was used as a control.

For adenoviral infection, MEFs were incubated in serum-free DMEM with 1 mg/ml BSA for approximately 9 hours. Adenoviruses were added to the culture medium and incubated for 24 hours. Adenoviruses encoding Rac^V12^ and Rac^N17^ were added at a multiplicity of infection of 900 and 100, respectively. Additionally, adenovirus encoding LacZ was used as a control. Adenovirus-infected MEFs were seeded onto fibronectin-coated hydrogels with DMEM containing 10% FBS.

In some experiments, adenovirus-infected cells were cultured on fibronectin-coated hydrogels for 24 hours with DMSO (vehicle, Sigma-Aldrich) or 0.5 μM latrunculin B (Calbiochem).

### Preparation of fibronectin-coated polyacrylamide hydrogels

The general protocol for fabricating stiffness-tunable polyacrylamide hydrogels has been previously described [27, 28]. Briefly, glass coverslip surfaces were etched uniformly with 0.1 M sodium hydroxide solution (Fisher), followed by the addition of (3-Aminopropyl)trimethoxysilane (APTMS; Sigma-Aldrich) to introduce amine (-NH2) groups to the glass surface. Reactive coverslips were then incubated with 0.5% glutaraldehyde solution (Sigma-Aldrich) to cross-link the APTMS and the polyacrylamide hydrogels. Hydrogels of varying stiffness were prepared using varying ratios of 40% acrylamide to 1% bis-acrylamide (Bio-Rad) in a mixed solution with water, ammonium persulfate (Sigma-Aldrich), TEMED (Bio-Rad), and acrylic acid N-hydroxysuccinimide ester (NHS, Sigma-Aldrich) in toluene (Sigma-Aldrich). Notably, the polyacrylamide hydrogels were functionalized using NHS solution to induce the conjugation of fibronectin (Sigma-Aldrich) to the polyacrylamide hydrogels. The elastic moduli of the hydrogels with varying stiffness were used for the experiment: low (2 to 4 kPa), medium (10 to 12 kPa), and high (20 to 24 kPa). The hydrogels were then coated with 3 μg/mL fibronectin and incubated overnight at 4°C. The fibronectin-coated hydrogels were extensively washed with DPBS to remove monoacrylamide. Unreactive cross-linkers were blocked with 1 mg/ml heat-inactivated BSA prior to cell seeding. Glass coverslips of different sizes and seeded cell densities were used for specific experiments: immunoblotting and Rac1 G-LISA assay (40 mm, 2 x 10^5^ cells), immunofluorescence and atomic force microscopy (12 mm, 2 x 10^4^ cells), and RT-qPCR (25 mm, 10^5^ cells).

### RNA isolation and real-time quantitative PCR (RT-qPCR)

MEFs cultured on polyacrylamide hydrogels were first washed twice with ice-cold DPBS and then collected using TRIzol reagent (Invitrogen). Total RNA was extracted according to the manufacturer’s instructions as described [27]. Briefly, TRIzol extracts were incubated with chloroform for 3 minutes at room temperature (RT) and then centrifuged at 15,000 rpm for 15 minutes at 4°C. Thereafter, the upper aqueous phase containing RNA was transferred to a new Eppendorf tube, and RNA was incubated with 100% isopropanol for 10 minutes at RT. Samples were then centrifuged at 15,000 rpm for 15 minutes at 4°C. The supernatant was discarded, and the RNA pellet was resuspended in 75% ethanol. The samples were centrifuged at 9,000 rpm for 5 minutes at 4°C. RNA was eluted with RNAse-free water and then incubated at 55 to 60°C for 10 minutes. Total RNA was reverse transcribed and analyzed by RT-qPCR as previously described. TaqMan assays were used for lamellipodin, cyclin A, cyclin D1, and GAPDH. The relative changes in mRNA expression for each target gene were determined by the delta-delta-Ct method using GAPDH as the reference gene.

### Protein extraction, immunoblotting, and immunostaining

Protein extraction was performed as follows: adherent MEFs on fibronectin-coated polyacrylamide hydrogels were first washed twice with ice-cold DPBS, and total cell lysates were then extracted by placing the hydrogel face-down on a predetermined volume of 5X sample buffer (250 mM Tris pH 6.8, 10% SDS, 50% glycerol, 0.02% bromophenol blue, and 10 mM 2-mercaptoethanol) and incubated for 2 minutes at RT, as previously described [2, 27]. Equal amounts of extracted protein were fractionated in 5 to 10% SDS-PAGE gels, and the fractionated proteins were transferred electrophoretically onto polyvinylidene difluoride membranes. These immunoblot membranes were probed with antibodies against lamellipodin (Santa Cruz Biotechnology), cyclin D1 (Santa Cruz Biotechnology), cyclin A [29], cyclin B1 (Santa Cruz Biotechnology), Rac1 (Santa Cruz Biotechnology), vinculin (Sigma-Aldrich), and GAPDH (Proteintech). Antibody signals were detected using enhanced chemiluminescence (Bio-Rad).

For immunostaining, MEFs cultured on polyacrylamide hydrogels were fixed in 3.7% formaldehyde for 1 hour, permeabilized with 0.4% Triton X-100 for 30 minutes, and then blocked with 2% BSA and 0.2% Triton X-100 for 1 hour at RT. To observe filamentous actin (F-actin), the cells were incubated with Alexa Fluor 488 Phalloidin (Invitrogen) for 2 hours at RT. Then, the cells were washed with DPBS containing 2% BSA and 0.2% Triton X-100 and then with distilled water before mounting for confocal microscopy. Fluorescence images of the cells were acquired with a 20X objective of a VisiTech VT-I iSIM confocal microscope system (VisiTech International).

### Atomic Force Microscopy (AFM)

AFM was used to measure intracellular stiffness as previously described [1, 2, 30]. Briefly, to measure intracellular stiffness, MEFs cultured on soft and stiff polyacrylamide hydrogels in phenol red-free DMEM with 10% FBS were indented with a silicon nitride cantilever (Bruker; spring constant, 0.06 N/m) with a conical AFM tip (40 nm in diameter). AFM in contact mode was applied to single adherent MEFs using a BioScope Catalyst AFM system (Bruker) mounted on a Nikon Eclipse TE 200 inverted microscope. To analyze the stiffness, the first 600 nm of horizontal tip deflection was fit with the Hertz model for a cone. 5 to 10 measurements of intracellular stiffness from each experiment were acquired near the periphery of each cell (8 to 10 cells per experimental condition). AFM curves were quantified and converted to Young’s modulus (stiffness) using AFM analysis software (NanoScope Analysis, Bruker).

### Statistical analysis

Data are presented as the mean + standard error of the mean deviation (SE) of the indicated number of independent experiments. The data were analyzed using Student’s t-test. The results with p-values lower than 0.05 (*), 0.01 (**), or 0.001 (***) were considered to be statistically significant.

## ACKNOWLEDGMENTS

This work was funded in part by the American Heart Association Career Development Award (18CDA34080415) and the State University of New York start-up funding to Y.B. The funders had no role in study design, data collection and analysis, decision to publish, or preparation of the manuscript.

## AUTHOR CONTRIBUTIONS

J.A.B. and Y.B. designed the study and performed experiments and data analysis. J.A.B., K.L., and Y.B. wrote the manuscript. All authors critically reviewed and edited the manuscript.

## DECLARATION OF INTERESTS

The authors declare no competing interests.

## Notes

### Competing Interest Statement

The authors have declared no competing interest.

